# Isoform-Specific Control of Adipose Thermogenesis by the miR-27 Family Reveals Antagonism within a Polycistronic miRNA Cluster

**DOI:** 10.1101/2025.07.21.666011

**Authors:** Devesh Kesharwani, Michele Karolak, Chad C. Doucette, Eric R. Loomis, Sophia E Adami-Sampson, Su Su, Victoria DeMambro, Clifford Rosen, Anne Harington, Larisa Ryzhova, Lucy Liaw, Aaron C. Brown

**Author notes:** Correspondence. Aaron C. Brown. Lead contact. Aaron C. Brown. A.C.B.: Conceptualization, Methodology, Formal analysis, Writing – Original Draft, Supervision, Funding acquisition. D.K.: Investigation, Formal analysis, Writing – Original Draft, Review & Editing. M.K.: Investigation, Formal analysis. C.C.D.: Investigation. E.R.L.: Investigation. S.S.: Investigation. V.D.: Investigation, Formal analysis. A.H.: Resources. L.R.: Resources. C.R.: Resources. L.L.: Resources, Review & Editing, Funding acquisition.

## Abstract

MicroRNAs (miRNAs) are frequently encoded within polycistronic clusters thought to function as coherent regulatory units, yet whether individual cluster members act cooperatively or exert distinct physiological effects remains unclear. miR-27a and miR-27b are closely related miRNAs located within separate paralogous miR-23/27/24 clusters, but their *in vivo* relationship is unresolved. Using isoform-specific knockout models generated by precise CRISPR-based excision of individual miRNA hairpins, we defined how miR-27a and miR-27b regulate adipose thermogenesis and systemic metabolism. High-fat feeding increased miR-27a/b expression and suppressed thermogenic gene programs in subcutaneous white adipose tissue. In primary beige and brown adipocytes, loss of either isoform produced modest activation of thermogenic programs, whereas combined deletion caused additive increases in *Ucp1* expression and mitochondrial DNA content. *Ex vivo* heat production was significantly elevated in double-knockout adipose explants relative to wild-type and single-knockout controls. *In vivo*, double-knockout mice exhibited increased energy expenditure and, under high-fat diet conditions, reduced adiposity with improved glucose homeostasis compared with wild-type and single-knockout mice. In contrast to metabolic dysfunction reported following whole-cluster deletion, selective miR-27 loss reveals a protective thermogenic program. These findings demonstrate that miRNA clusters need not act as coherent regulatory modules and identify miR-27a/b as a cooperative thermogenic checkpoint with therapeutic implications.

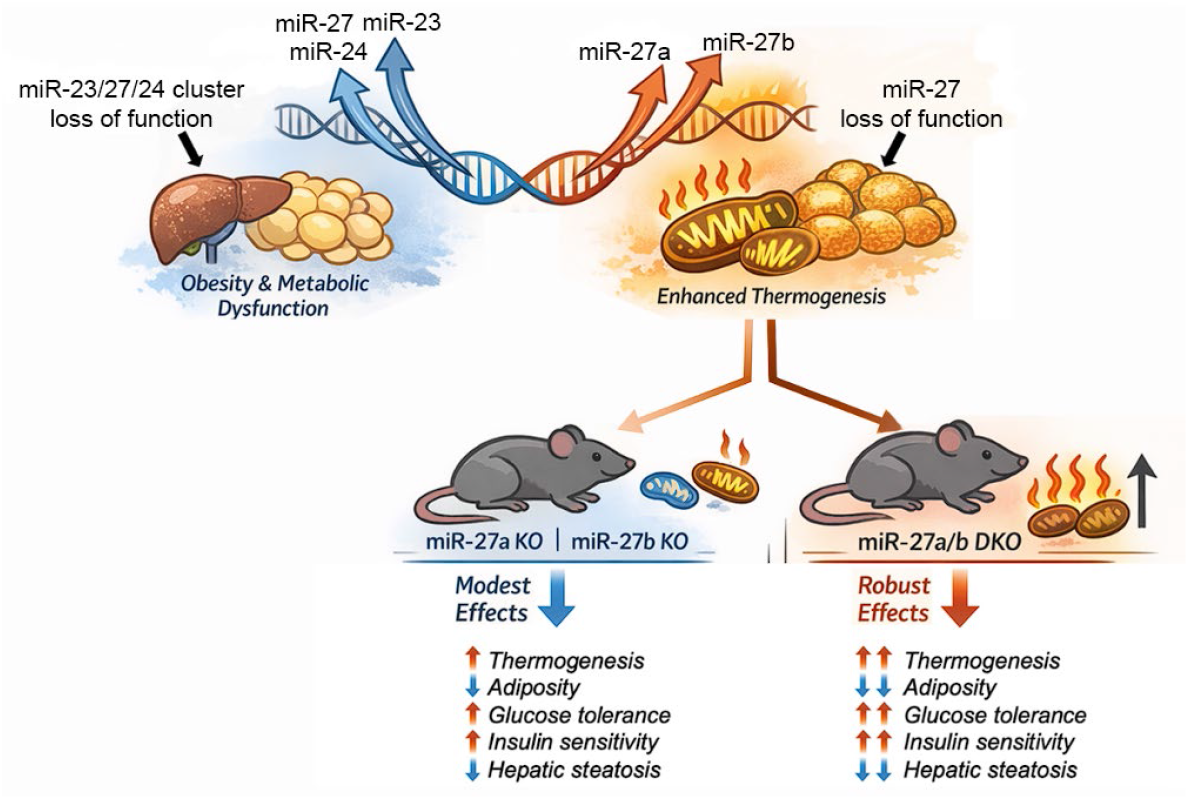

* CRISPR isoform-specific knockouts isolate miR-27a/b contributions to metabolism.

*Dual miR-27a/b loss drives cooperative thermogenic activation.

*Double knockouts enhance energy expenditure and metabolic fitness in diet-induced obesity.

*miR-27a/b deletion reveals that clustered miRNAs can exert non-coherent regulatory roles.

## Introduction

Obesity has become a major global health challenge, driving rising rates of chronic disease, diminishing quality of life, and placing substantial burdens on healthcare systems. The rising prevalence of obesity is closely linked to increased risk of metabolic-associated disorders, including type 2 diabetes, cardiovascular disease, and metabolic dysfunction-associated steatotic liver disease (MASLD), and is further associated with heightened susceptibility to several types of cancer^1-4^. The pathogenesis of obesity is multifactorial, driven by a complex interplay of genetic predisposition, environmental influences, and epigenetic regulators. Together, these factors disrupt the tightly regulated balance of energy homeostasis by influencing appetite control, nutrient absorption, and energy expenditure^5, 6^.

Adipose tissue is a central regulator of systemic energy homeostasis, with white adipose tissue (WAT) serving as a primary depot for energy storage and brown adipose tissue (BAT) functioning as a site for energy dissipation through thermogenesis^7^. In obesity, excessive expansion and dysfunction of WAT contribute to metabolic impairment. Enhancing energy expenditure by activating thermogenesis within adipose depots has therefore emerged as a promising therapeutic strategy^8^. Central to this process is uncoupling protein 1 (UCP1), which enables the conversion of chemical energy into heat in brown adipocytes and in inducible beige adipocytes within subcutaneous WAT^9^. Consequently, defining the molecular pathways that regulate *Ucp1* expression and promote the browning of WAT will provide a foundation for the development of new interventions to counteract obesity

MicroRNAs (miRNAs) are critical post-transcriptional regulators of gene expression and play essential roles in adipogenesis, thermogenesis, and systemic metabolic regulation^10, 11^. Several miRNAs, including miR-328, miR-378, miR-455, miR-30, and miR-32, promote brown adipocyte differentiation and thermogenic activation^12-16^. In contrast, miR-34a, miR-27a, miR-27b, miR-155, and miR-133 inhibit brown adipogenesis^17-20^. Among these, the miR-27 family, which includes miR-27a and miR-27b, has emerged as a central negative regulator of adipogenesis and thermogenesis^21, 22^. Both isoforms are downregulated during adipocyte differentiation and inhibit thermogenic programming by repressing transcriptional coactivators and regulatory factors. Inhibition of miR-27 in cultured beige and brown adipocytes increases expression of thermogenic genes, including *Ucp1, Prdm16*, and *Pgc1α*, while overexpression of miR-27 suppresses their expression^22^. Mechanistically, miR-27b directly targets the 3′-untranslated regions (3′-UTRs) of *Pparα, Pparγ, Prdm16, Pgc1β* and *Creb1*, thereby restraining adipocyte differentiation and thermogenic activity^22-24^. Levels of miR-27a and miR-27b are elevated in WAT from high-fat diet (HFD)-induced obese mice, supporting their role in suppressing thermogenesis and contributing to obesity^23, 24^. Conversely, antagomir-mediated inhibition of miR-27b was shown to enhance thermogenic gene expression of epididymal fat and confer resistance to obesity^25^. However, antagomir-based approaches are subject to off-target effects, transient suppression, and dose-dependent variability, which complicates the interpretation of their physiological impact. Despite these insights, the *in vivo* consequences of complete genetic loss of miR-27a, miR-27b, or both isoforms have not been fully characterized. As a result, the extent to which each isoform contributes to energy homeostasis and thermogenic regulation remains unclear.

Beyond antagomir-mediated inhibition, several groups have used genetic approaches to interrogate the miR-23/27/24 clusters, in which miR-27a and miR-27b are embedded as part of conserved tricistronic loci. Global deletion of the miR-23b/27b/24-1 cluster resulted in impaired glucose tolerance and systemic metabolic dysfunction, with prominent alterations in hepatic metabolic pathways^26^. In a conditional model, myeloid-specific deletion of both the miR-23a/27a/24-2 and miR-23b/27b/24-1 clusters led to exacerbated obesity-associated inflammation and worsening insulin resistance^27^. Together, these cluster-level genetic studies demonstrate that loss of the miR-23/27/24 loci adversely impacts metabolic physiology; however, because all cluster constituents were simultaneously deleted, the specific contributions of individual miR-27 isoforms and their intrinsic roles in adipose tissue thermogenic regulation remain unresolved.

To address this gap, we examined the distinct and overlapping roles of miR-27a and miR-27b in regulating thermogenesis and obesity susceptibility using newly generated genetic knockout models that provide a more rigorous and physiologically relevant framework for defining isoform-specific functions *in vivo*. Although miR-27a and miR-27b are closely related, subtle sequence and regulatory differences between the two isoforms may produce non-identical biological effects. Across cellular and tissue assays, deletion of either miR-27a or miR-27b produced modest thermogenic activation, whereas combined loss of both isoforms resulted in additive increases in *Ucp1* expression, mitochondrial DNA content, and adipose tissue thermogenic output relative to wild-type and single-knockout controls. Under high-fat diet conditions, all knockout genotypes exhibited reduced adiposity compared with wild-type mice without evidence of cooperativity. In functional metabolic phenotyping, double-knockout mice displayed increased energy expenditure versus wild-type controls and improved fasting glycemia, glucose tolerance, and insulin sensitivity compared with both wild-type and single-knockout mice.

Together, these results demonstrate that simultaneous loss of miR-27a and miR-27b is required to fully activate adipocyte thermogenesis and achieve maximal metabolic protection, and indicate that whole-cluster deletion likely masks the beneficial effects of miR-27 loss by concurrently removing miRNAs that support metabolic homeostasis. More broadly, this work reveals how coordinated repression by closely related miRNAs can shape thermogenic fate decisions, advancing our understanding of the regulatory logic governing adipose tissue plasticity. In doing so, this study identifies the miR-27 family as a central suppressor of thermogenic programming and underscores its potential as a therapeutic target for obesity.

## Results

### HFD Drives Metabolic Dysfunction with Altered miR-27a/b and Thermogenic Gene Expression

To determine how obesity influences adipose tissue physiology, we employed a diet-induced obesity model using 3-week-old male C57BL/6J mice (Fig. 1A). Mice were fed either a standard control or HFD for 16 weeks. HFD-fed mice exhibited a significant increase in body weight compared with chow controls (Fig. 1B) and developed hyperglycemia, as evidenced by elevated fasting glucose levels (Fig. 1C). Glucose tolerance tests (GTT) and insulin tolerance tests (ITT) further confirmed impaired glucose clearance and reduced insulin sensitivity in HFD-fed mice (Fig. 1D-E). After 16 weeks, mice were euthanized and adipose depot weights were assessed, including brown adipose tissue (BAT), subcutaneous WAT (inguinal), and gonadal WAT (Fig. 1F-G). HFD feeding resulted in a robust increase in adiposity, with inguinal WAT mass increasing three-fold and gonadal WAT mass increasing to nearly five-fold compared with controls.

**Figure 1.**
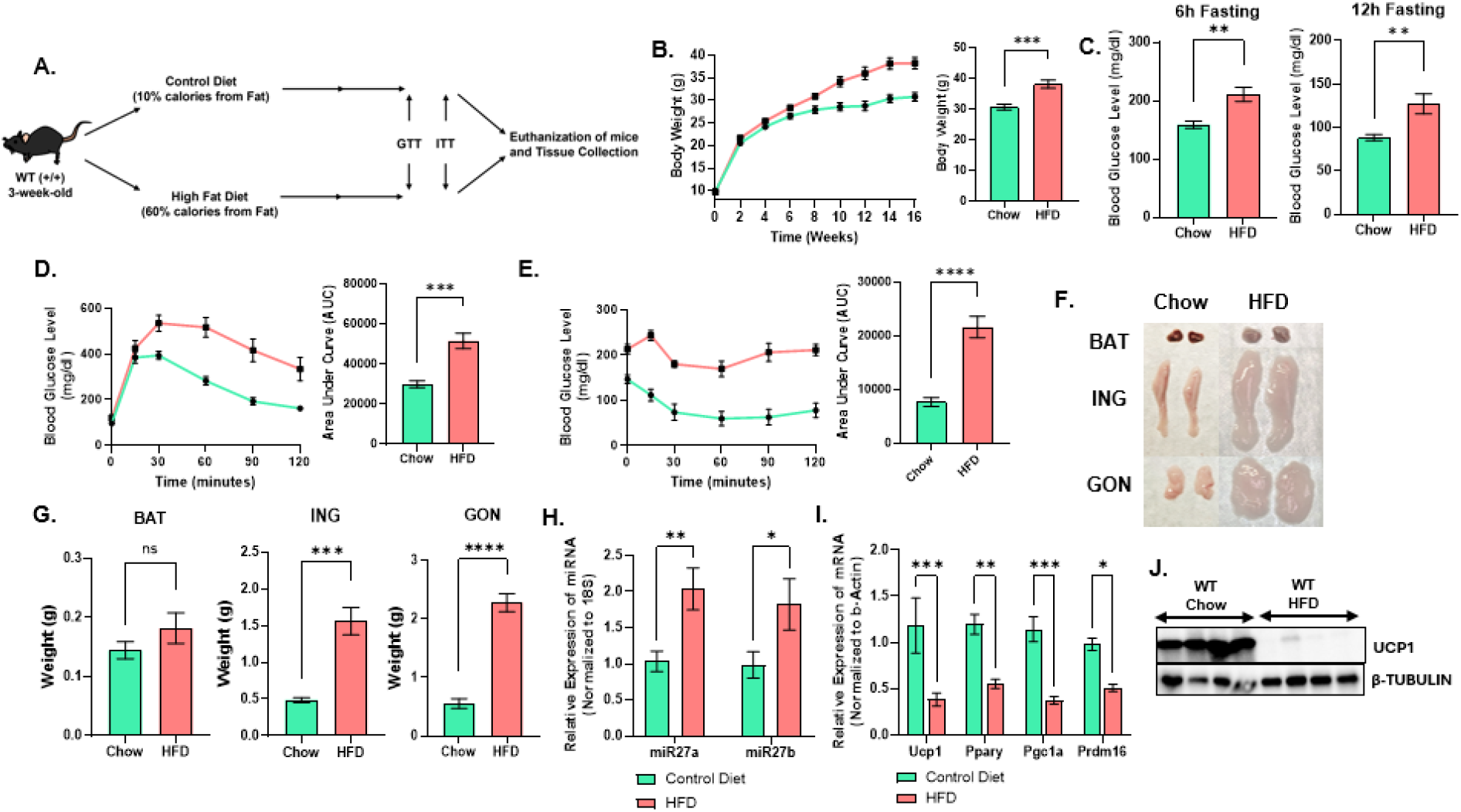
HFD drives metabolic dysfunction with altered miR-27a/b and thermogenic gene expression. **(A)** Schematic of the diet-induced obesity protocol used in C57BL/6J wild-type mice. **(B)** Body weight trajectory during chow or HFD feeding (n = 7). **(C)** Fasting blood glucose levels after 15 weeks of chow or HFD. **(D)** Glucose tolerance test and **(E)** insulin tolerance test performed after 16 weeks on chow or HFD (n = 7). **(F)** Representative appearance of brown adipose tissue (BAT), inguinal white adipose tissue (ING), and gonadal white adipose tissue (GON) at the end of the study. **(G)** Weight of BAT, ING, and GON depots at euthanasia (n = 7). **(H)** TaqMan-based quantification of miR-27a and miR-27b expression in ING adipose tissue from chow and HFD mice. **(I)** qRT-PCR analysis of thermogenic and adipogenic gene expression in ING adipose tissue from chow and HFD mice. **(J)** Western blot analysis of UCP1 protein levels in ING adipose tissue from chow and HFD mice. Data are presented as mean ± SEM. Statistical significance is indicated as *p < 0.05, **p < 0.01, ***p < 0.001, and ****p < 0.0001.

To assess the impact of diet-induced obesity on browning (beige adipocyte induction) and thermogenic programming of subcutaneous WAT, we measured miR-27a/b expression and thermogenic gene markers in the inguinal adipose depot from mice fed either a standard control diet or HFD. Building on previous work showing that miR-27a/b are suppressed in subcutaneous WAT during cold-induced thermogenic activation^22^, miR-27a and miR-27b were significantly upregulated in inguinal WAT of HFD-fed mice compared with controls (Fig. 1H). This increase was accompanied by a pronounced reduction in key thermogenic and adipogenic regulators, including *Ucp1, Pparγ, Pgc1α*, and *Prdm16* (Fig. 1I). Western blot analysis similarly revealed decreased UCP1 protein levels in inguinal WAT of HFD-fed mice (Fig. 1J). The inverse relationship between miR-27a/b expression and thermogenic gene programs suggests that elevated miR-27 family members contribute to suppression of adipose tissue thermogenic capacity and participate in adipose tissue remodeling during obesity development.

### Generation and Validation of Novel miR-27a and miR-27b Knockout Mice

miR-27a and miR-27b share an identical seed sequence, a key determinant of mRNA target recognition, and differ by only a single nucleotide in their mature forms (Fig. 2A), suggesting that they may regulate overlapping mRNA targets and compensate for one another. These miRNAs reside within the conserved and tightly organized miR-23a/27a/24-2 and miR-23b/27b/24-1 genomic clusters on mouse chromosomes 8 and 13, respectively, where the three miRNAs are initially transcribed as a single tricistronic primary transcript (pri-miRNA). Although global and conditional knockout mice targeting the miR-23a∼27a∼24-2 and miR-23b∼27b∼24-1 clusters have been generated and used to interrogate hematopoiesis^28, 29^, skeletal and cartilage biology^30, 31^, and whole-body glucose metabolism^26, 27^, the specific functional impact of deleting miR-27a and miR-27b individually within their respective clusters in relation to systemic metabolism and adipose biology, has not been addressed.

**Figure 2.**
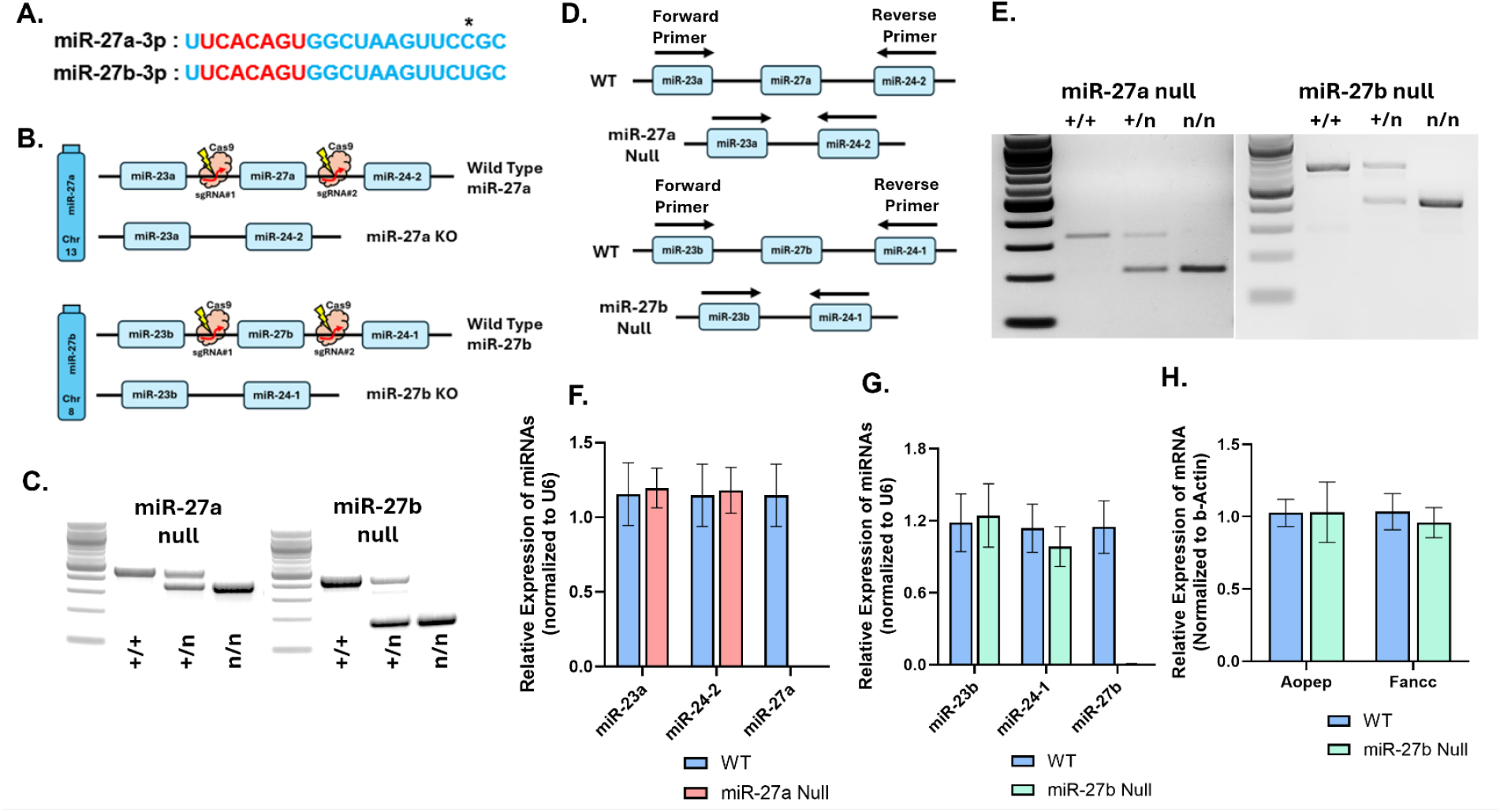
Generation and validation of novel miR-27a and miR-27b knockout mice. (A) Mature miRNA sequences of miR-27a and miR-27b highlighting their shared seed sequence and single-nucleotide difference. (B) Schematic of the CRISPR/Cas9 strategy used to generate miR-27a and miR-27b knockout mice, including guide RNA targeting sites. (C) Representative PCR genotyping showing wild-type (+/+), heterozygous (+/n), and homozygous null (n/n) alleles for miR-27a and miR-27b. (D) Strategy used to confirm that the tricistronic miR-23/miR-27/miR-24 clusters remain transcriptionally active after deletion of miR-27a or miR-27b, using primer pairs spanning miR-23a to miR-24-2 or miR-23b to miR-24-1, respectively. (E) Agarose gel images of RT-PCR products generated from RNA isolated from inguinal WAT of wild-type, heterozygous, and knockout mice, detecting amplification of the tricistronic primary miRNA (pri-miRNA) cluster transcript spanning flanking miR-23 and miR-24 loci. (F) qPCR analysis of miR-23a, miR-24-2, and miR-27a expression in inguinal adipose tissue from wild-type and miR-27a-null mice (n = 8). (G) qPCR analysis of miR-23b, miR-24-1, and miR-27b expression in inguinal adipose tissue from wild-type and miR-27b-null mice (n = 8). (H) qPCR of *Aopep* and *Fancc* mRNA expression in inguinal adipose tissue from wild-type and miR-27b-null mice showing stable host-gene expression after deletion of intronic miR-27b (n = 8). Data are shown as mean ± SEM.

To investigate the individual and combined roles of miR-27a and miR-27b in adipose tissue thermogenesis and adipogenesis, we generated global miR-27a and miR-27b knockout mice using a CRISPR/Cas9-mediated deletion strategy (Fig. 2B). Guide RNAs (gRNAs) with high predicted specificity and efficiency were designed to target each locus, and single-cell C57BL/6J embryos were microinjected with the CRISPR/Cas9 complex before transfer into pseudo-pregnant surrogate females. Resulting heterozygous pups were used as founders and backcrossed to the C57BL/6J background for more than ten generations to establish stable miR-27a and miR-27b knockout lines. Homozygous deletions were confirmed by PCR-based genotyping (Fig. 2C).

To ensure that deletion of miR-27a or miR-27b did not disrupt transcription of adjacent miRNAs within their respective tri-miRNA clusters, we designed RT-PCR primers flanking the deleted miRNA loci to amplify the pri-miRNAs spanning miR-23 to miR-24 (miR-23a forward and miR-24-2 reverse for the miR-27a cluster, and miR-23b forward and miR-24-1 reverse for the miR-27b cluster) (Fig. 2D). RT-PCR analysis was performed on RNA isolated from inguinal adipose tissue, demonstrating continued amplification of the pri-miRNA products in wild-type, heterozygous, and miR-27 knockout mice (Fig. 2E), confirming that cluster transcription remained intact despite selective deletion of miR-27. Consistent with this, quantitative PCR (qPCR) analysis showed comparable expression of the adjacent mature miRNAs miR-23a and miR-24-2 in miR-27a knockout mice, and miR-23b and miR-24-1 in miR-27b knockout mice, relative to wild-type controls (Fig. 2F-G). These results demonstrate that targeted excision of individual miR-27 hairpins did not impair transcription or processing of the surrounding tricistronic pri-miRNA clusters, allowing phenotypic changes observed in these mice to be attributed specifically to loss of miR-27a or miR-27b rather than disruption of cluster architecture.

Because miR-27a and miR-27b reside in distinct genomic contexts, with miR-27a located in an intergenic cluster on chromosome 8 and miR-27b located within the last intron of the *Aopep* gene on chromosome 13 in a compact gene-dense region adjacent to *Fancc*, we next evaluated whether deletion of the intronic miR-27b hairpin altered expression of nearby host genes. mRNA levels of *Aopep* and *Fancc* in inguinal WAT were measured using exon-spanning primers with qPCR (Fig. 2H). Both transcripts exhibited stable expression in miR-27b knockout mice compared with wild-type controls, indicating that the miR-27b deletion did not disrupt regulatory elements required for proper host-gene expression.

Breeding viability and progeny distributions were assessed across all genotypes to determine whether loss of miR-27a, miR-27b, or both miRNAs affected reproductive fitness (Supplementary Table 1). Comparable numbers of breeder pairs were established for wild-type, miR-27a null, miR-27b null, and DKO lines, with similar conception rates observed across groups. Litter sex ratios did not deviate from the expected 1:1 distribution for any genotype and were not significantly different from wild-type controls (p > 0.89 for all comparisons; Suppl. Table 1A). Although fertility rates and mean litter sizes were modestly reduced in knockout lines, particularly in the DKO genotype (Suppl. Table 1B), all lines were viable and capable of sustaining colony maintenance and experimental cohort generation.

Collectively, these findings demonstrate that miR-27a and miR-27b can be specifically and effectively ablated without altering expression of neighboring miRNAs and local host genes, or fundamental breeding viability, enabling direct assessment of their individual and combined contributions to metabolic regulation.

### Combined deletion of miR-27a and miR-27b enhances thermogenic function

To determine how single and combined loss of miR-27a and miR-27b influences adipocyte thermogenesis, we isolated the stromal vascular fraction (SVF) from inguinal WAT of wild-type (WT), miR-27a-null, miR-27b-null, and DKO mice and differentiated these cells into beige adipocytes. TaqMan-based qPCR confirmed complete loss of miR-27a and miR-27b expression in adipocytes from the corresponding knockout lines (Fig. 3A). Multiple thermogenic-associated genes showed broad upregulation in beige adipocytes derived from miR-27a-, miR-27b-, and DKO mice relative to WT; however, apart from *Dio2* (which appeared additive), their expression did not appear to be additively or synergistically increased in the DKO (Fig. 3B-D). In contrast, deletion of both miRNAs produced a markedly stronger induction of *Ucp1* expression compared with WT and either single knockout (Fig. 3E). Notably, *Ucp1* expression in DKO beige adipocytes showed an additive increase relative to the single knockouts, suggesting a complementary contribution of miR-27a and miR-27b toward suppression of *Ucp1* expression under basal conditions.

**Figure 3.**
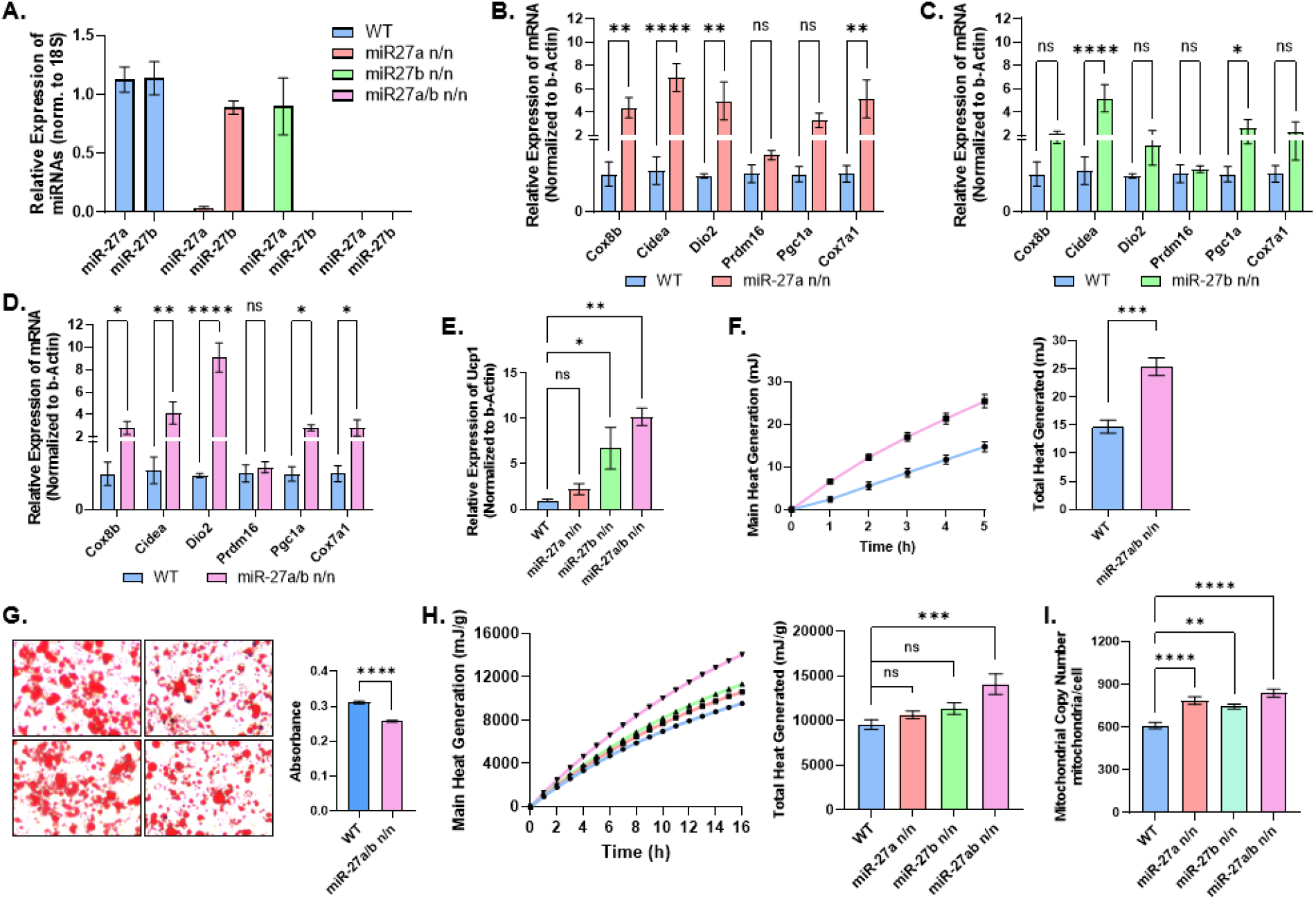
Combined deletion of miR-27a and miR-27b enhances thermogenic activity in adipocytes and adipose tissue. **(A)** TaqMan qPCR analysis of miR-27a and miR-27b expression in differentiated beige adipocytes derived from the stromal vascular fraction (SVF) of inguinal adipose tissue from WT, miR-27a KO, miR-27b KO, and miR-27a/b DKO mice (n = 4). **(B-E)** qPCR analysis of thermogenic gene expression, including *Ucp1*, in differentiated beige adipocytes from WT, miR-27a KO, miR-27b KO, and DKO mice on day 10 of differentiation (n = 3). **(F)** Isothermal microcalorimetry of heat production from cultured beige adipocytes differentiated from SVF isolated from WT and DKO inguinal adipose tissue (n = 5). Left panel shows cumulative heat output over time; right panel shows total heat production quantified from time-course measurements. **(G)** Oil Red O staining of cultured beige adipocytes from WT and DKO mice. Left panel shows representative images; right panel shows quantification of lipid accumulation. **(H)** Isothermal microcalorimetry of heat production from inguinal adipose tissue explants isolated from 8-week-old WT, miR-27a KO, miR-27b KO, and DKO mice (n = 7). Left panel shows cumulative heat output over time; right panel shows quantification of total heat production. **(I)** Mitochondrial DNA copy number analysis in cultured beige adipocytes isolated from WT, miR-27a KO, miR-27b KO, and DKO mice (n = 4). Data are presented as mean ± SEM. Statistical significance was determined as *p < 0.05, **p < 0.01, ***p < 0.001, ****p < 0.0001.

To directly measure functional thermogenic output, we first performed isothermal microcalorimetry on cultured beige adipocytes. This analysis revealed significantly higher heat production in DKO beige adipocytes compared with WT cells (Fig. 3F). This increase was also associated with reduced lipid content in DKO adipocytes, possibly reflecting enhanced lipid utilization, despite the fact that *PPARγ*, an adipogenic regulator suppressed by miR-27, would normally be expected to promote increased adipogenesis and lipid storage (Fig. 3G)^32^. To determine whether this phenotype persisted *in vivo*, microcalorimetry was performed on explants of inguinal WAT isolated from WT, miR-27a, miR-27b, and DKO mice reared at room temperature. Consistent with the *in vitro* findings, inguinal WAT from DKO mice exhibited significantly increased heat output compared with WT and single knockouts (Fig. 3H). Because enhanced heat generation in thermogenic adipocytes often correlates with increased mitochondrial abundance, mitochondrial DNA copy number was quantified in isolated inguinal-derived beige adipocytes from each genotype. DKO adipocytes displayed a significant increase in mitochondrial DNA copy number relative to WT and single single knockouts (Fig. 3I). Parallel analysis of brown adipocytes revealed similar patterns, including increased heat generation in cultured DKO brown adipocytes versus wild-type controls, as well as elevated thermogenic output from DKO interscapular brown adipose tissue explants and increased mitochondrial DNA copy number in isolated interscapular-derived brown adipocytes, compared with wild-type and single-knockout controls (Suppl. Fig. 1A-C).

Together, these results indicate that while single deletion of miR-27a or miR-27b produces modest thermogenic activation, the combined loss of both miRNAs yields a substantially stronger response. The DKO showed the greatest increases in *Ucp1*, thermogenic output, mitochondrial abundance, and lipid utilization, supporting a cooperative role for miR-27a and miR-27b in limiting adipose tissue thermogenic capacity.

### Deletion of miR-27a/b Increases Whole-Body Energy Expenditure

To assess whether deletion of miR-27a/b alters whole-body metabolism, age-matched WT and DKO male mice were singly housed at thermoneutrality (29.5°C) for 6 days. Thermoneutral housing minimizes the cold-induced thermogenesis that occurs at standard room temperature, reducing sympathetic activation and between-animal variability before metabolic cage analysis. Mice were maintained at thermoneutrality for an additional 4 days in metabolic cages, then sequentially exposed to room temperature (22°C) for 3 days and cold (6.5°C) for 3 days (Fig. 4A). Throughout this period, oxygen consumption (VO_2_), carbon dioxide production (VCO_2_), respiratory exchange ratio (RER), total energy expenditure (EE), resting energy expenditure (REE), physical activity, food intake, and water intake were continuously monitored.

**Figure 4.**
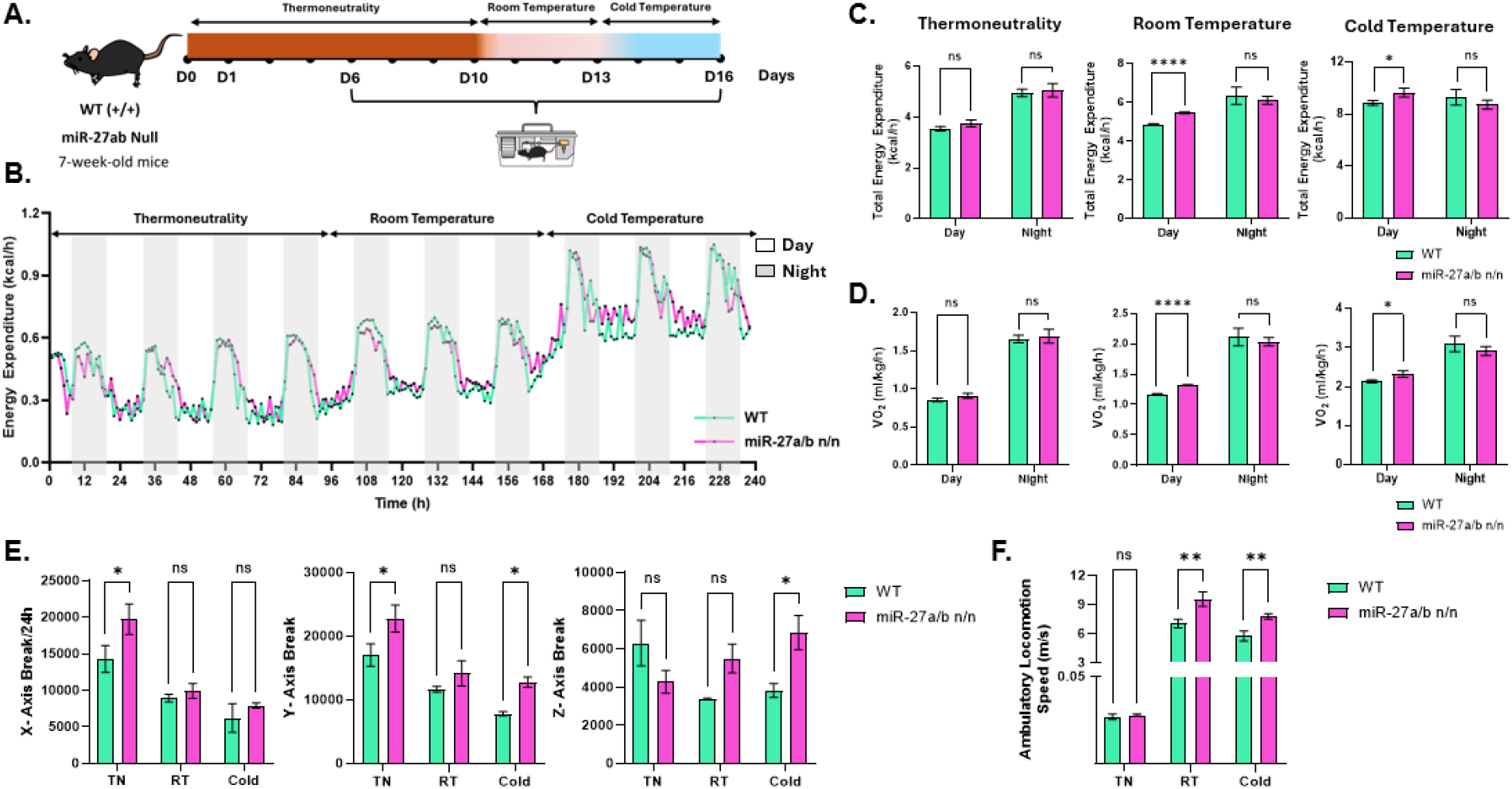
Deletion of miR-27a/b Increases Whole-Body Energy Expenditure. **(A)** Schematic of the metabolic cage temperature-challenge protocol. Seven-week-old male wild-type (WT) and miR-27a/b double-knockout (DKO) mice were acclimated at thermoneutral temperature (28 °C) for 6 days in an incubator, followed by transfer to metabolic cages. Mice remained at thermoneutrality for an additional 4 days for baseline measurements, then were sequentially exposed to room temperature (22 °C) for 3 days and cold temperature (6.5 °C) for 3 days (n = 3 per group). **(B)** Continuous measurement of total energy expenditure over time across all temperature conditions. **(C)** Quantification of mean total energy expenditure and **(D)** oxygen consumption (VO_2_) at thermoneutral, room temperature, and cold exposure. **(E)** Locomotor activity measured as axis beam breaks and (F) ambulatory movement across temperature conditions. Data are presented as mean ± SEM. Statistical significance is indicated as *p < 0.05, **p < 0.01, and ****p < 0.0001.

DKO mice exhibited significantly higher daytime energy expenditure and oxygen consumption at both room and cold temperatures compared with WT controls, whereas nighttime values remained comparable between genotypes (Fig. 4B-D). Similarly, DKO mice displayed increased carbon dioxide production and elevated resting energy expenditure during the lowest 30-minute activity period (REE_30) under both room and cold conditions during the day and under thermoneutral conditions at night (Suppl. Fig. 2A-B).

Food and water intake did not differ between genotypes at any temperature tested (not shown). However, physical activity analysis revealed condition-dependent differences between genotypes. DKO mice displayed higher locomotor and ambulatory activity than WT at some temperature conditions, and these differences appeared sporadically across conditions without implying any predictable pattern. Importantly, these fluctuations are unlikely to impact overall energy expenditure, as activity was not significantly different between groups at room temperature, and the DKO increase in energy expenditure occurred during the naturally low-activity daytime period (Fig. 4E-F).

Collectively, these findings indicate that miR-27a/b deficiency enhances oxygen consumption and energy expenditure, particularly under conditions that permit or increase thermogenic demand, including room temperature and cold exposure.

### Deletion of miR-27a/b Enhances Browning Capacity and Improves Metabolic Health in HFD-Induced Obesity

Given the inverse pattern between miR-27a/b expression and multiple thermogenic readouts, including higher *Ucp1* levels, increased heat production in adipocytes and adipose tissue, and elevated whole-body energy expenditure, we hypothesized that loss of miR-27a/b would enhance browning of adipose tissues and provide protection against diet-induced obesity. To test this, 3-week-old male WT, miR-27a-null, miR-27b-null, and DKO mice were placed on control or HFD for 16 weeks (Fig. 5A). Glucose and insulin tolerance tests (GTT and ITT) showed visibly improved glucose control and insulin sensitivity in all knockout groups, with the DKO displaying the most pronounced shift relative to WT (Fig. 5B-C). This included a significant reduction in the area under the curve (AUC), indicating a measurable improvement in glucose and insulin handling compared with the single knockouts.

**Figure 5.**
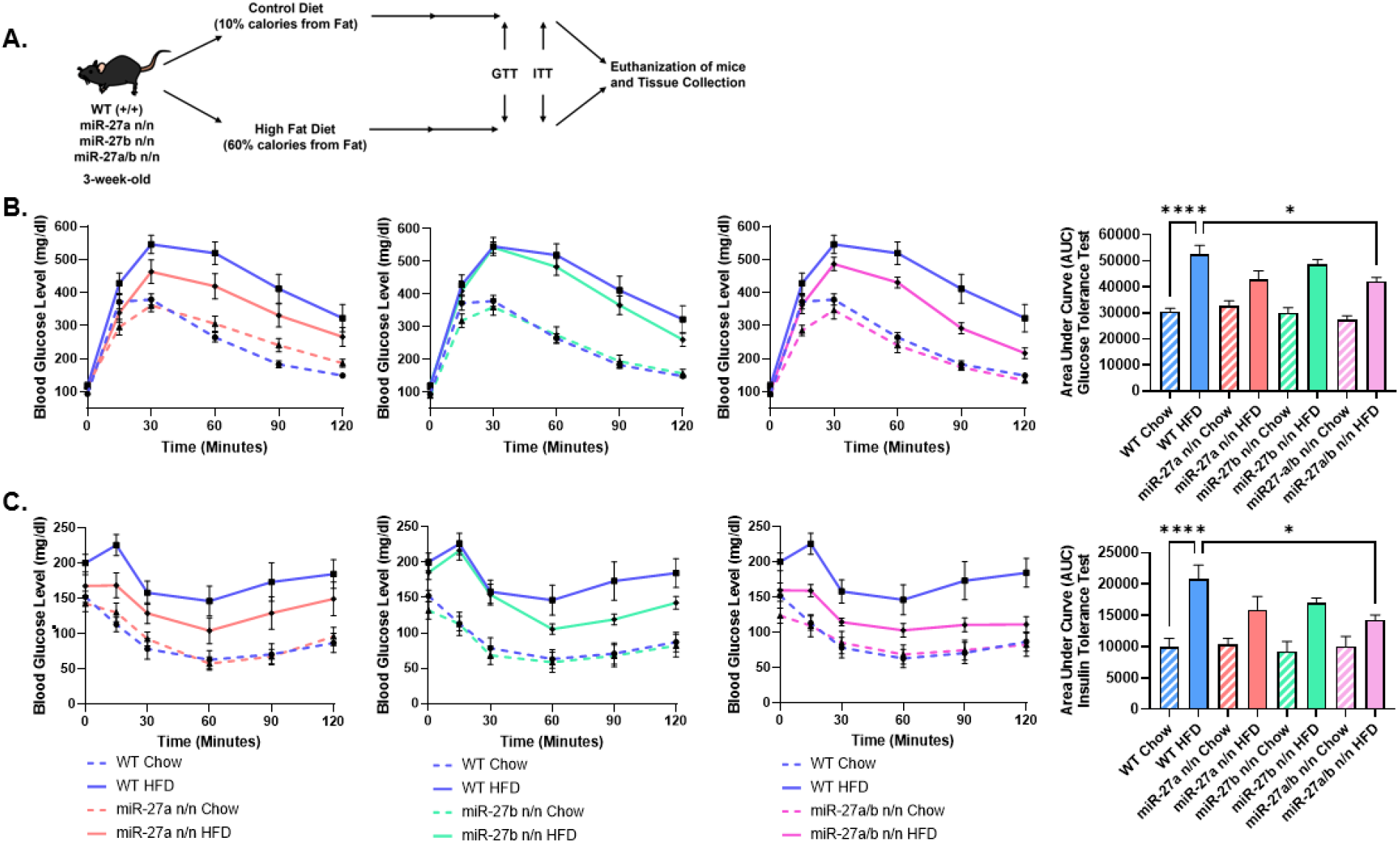
Deletion of miR-27a/b improves metabolic health under high-fat diet conditions. **(A)** Schematic of the diet-induced obesity protocol used for wWT, miR-27a KO, miR-27b KO, and miR-27a/b DKO male mice. **(B)** Glucose tolerance tests (GTT) performed after a 12-h fast in chow-or HFD-fed WT, miR-27a KO, miR-27b KO, and DKO mice (n = 9 per group). Blood glucose was measured over a 60-min period following glucose administration. Time-course data are shown with corresponding area-under-the-curve (AUC) quantification presented to the right. **(C)** Insulin tolerance tests (ITT) performed after a 6-h fast in chow- or HFD-fed WT, miR-27a KO, miR-27b KO, and DKO mice (n = 9 per group). Blood glucose was monitored for 60 min following insulin injection. Time-course data are shown with corresponding AUC quantification presented to the right. Data are presented as mean ± SEM. Statistical significance is indicated as *p < 0.05, **p < 0.01, ***p < 0.001, and ****p < 0.0001.

Male mice lacking miR-27b and the DKO exhibited significantly reduced body-weight gain under HFD conditions compared with WT (Fig. 6A, Suppl. Fig. 3). However, only the DKO showed a significant reduction in fasting blood glucose relative to WT HFD mice (Fig. 6B), indicating improved glucose homeostasis. After the 16-week diet, mice were euthanized, and brown, subcutaneous (inguinal), and gonadal adipose depots were dissected and weighed. Both inguinal and gonadal WAT depots from single knockouts and the DKO displayed significantly reduced fat mass, consistent with decreased lipid accumulation and attenuated adiposity (Fig. 6C-E). Hematoxylin and eosin (H&E) staining of BAT, inguinal WAT, and gonadal WAT revealed a pronounced reduction in lipid droplet size and overall lipid content across all adipose depots in DKO mice following HFD exposure (Fig. 6F). Notably, even under chow-fed conditions, all three depots in DKO mice contained a higher abundance of smaller, dense, multilocular adipocytes compared with WT, consistent with an enhanced basal browning phenotype.

**Figure 6.**
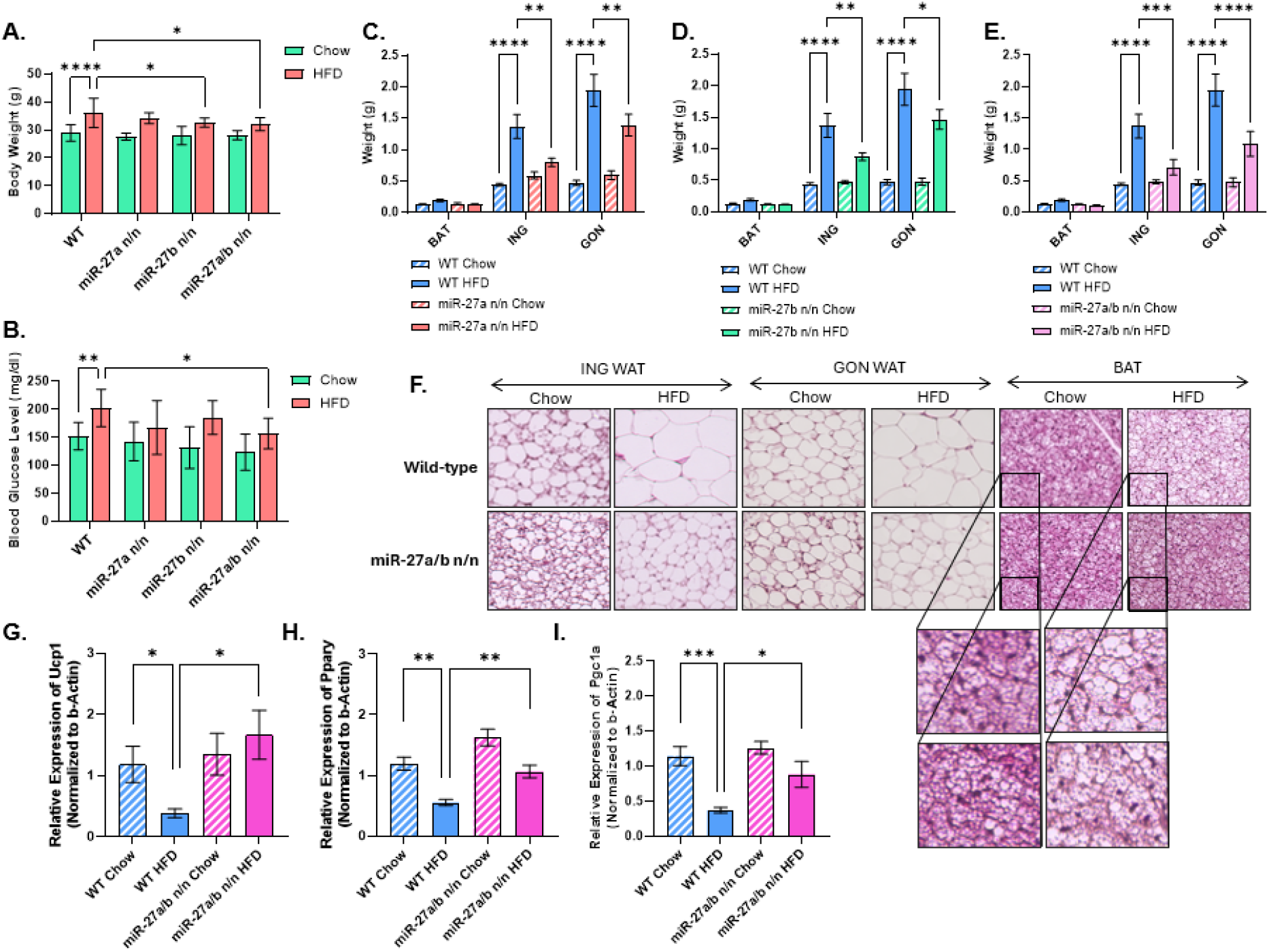
Combined deletion of miR-27a and miR-27b enhances adipose browning under high-fat diet conditions. **(A)** Body weight of WT, miR-27a KO, miR-27b KO, and miR-27a/b DKO male mice following 16 weeks of high-fat diet (HFD) feeding (n = 9 per group). **(B)** Fasting blood glucose levels measured after a 6-h fast at the conclusion of the HFD intervention (n = 9 per group). **(C-E)** Weights of brown adipose tissue (BAT; C), inguinal subcutaneous white adipose tissue (ING; D), and gonadal white adipose tissue (GON; E) collected from WT, miR-27a KO, miR-27b KO, and DKO mice maintained on either standard chow or HFD for 16 weeks (n = 9 per group). **(F)** Representative hematoxylin and eosin (H&E) staining of BAT, ING, and GON from WT and DKO mice following chow or HFD feeding, illustrating adipocyte morphology and lipid content. **(G-I)** Quantitative RT-PCR analysis of thermogenic and adipogenic gene expression in ING adipose tissue from WT and DKO mice, including *Ucp1* (G), *Pparγ* (H), and *Pgc1α* (I) (n = 7 per group). Data are presented as mean ± SEM. Statistical significance is indicated as *p < 0.05, **p < 0.01, ***p < 0.001, and ****p < 0.0001.

Gene expression analysis of inguinal adipose tissue revealed significant upregulation of key thermogenic markers, including *Ucp1* and *Pgc1α*, in HFD-fed DKO mice compared with WT HFD mice, along with increased expression of *Pparγ*, which is associated with adipocyte differentiation and beige adipocyte recruitment (Fig. 6F-H). These findings indicate that combined deletion of miR-27a and miR-27b enhances thermogenic gene expression and promotes WAT browning even under obesogenic conditions. Taken together, these results support a cooperative role for miR-27a and miR-27b in suppressing thermogenesis and promoting adiposity during HFD exposure, with dual deletion providing greater metabolic protection and browning capacity than loss of either miRNA alone.

## Discussion

Obesity arises from impaired energy balance, diminished thermogenic capacity, and reduced metabolic flexibility. Adipose tissue plasticity, particularly the ability of WAT to initiate thermogenic gene programs, is increasingly recognized as a central determinant of metabolic health^7, 9, 33^. In this study, we identify miR-27a and miR-27b as cooperative post-transcriptional repressors of adipocyte thermogenesis.

HFD feeding increased miR-27a/b expression while suppressing thermogenic gene activation in inguinal WAT, consistent with prior mouse studies demonstrating that miR-27 family members repress key regulators of adipocyte differentiation and thermogenic programming, including *Pparγ, Creb1*, and *Prdm16*^22-24, 32^. Similar inhibitory effects have been reported in human multipotent adipose-derived stem cells (hMADS), where miR-27b suppresses both PPARγ and CEBPα, limiting lipid accumulation and late-stage adipogenic gene expression^34^. In addition, miR-27a and miR-27b modulate mitochondrial integrity through prohibitin (PHB), a mitochondrial scaffold protein essential for cristae organization that is directly targeted by miR-27 and whose repression impairs adipocyte differentiation and mitochondrial function in hMADS^18^. Antagomir studies have shown that inhibition of miR-27a improves glucose uptake and insulin sensitivity in adipocytes and in HFD-fed mice^24^. Separately, Yu et al. demonstrated that antagomir-mediated inhibition of miR-27b enhances browning of epididymal adipose tissue and improves metabolic parameters in obese mice^25^. However, because these pharmacological approaches do not distinguish isoform-specific contributions or reveal potential functional cooperation, the extent to which miR-27a and miR-27b act independently or redundantly *in vivo* has remained unresolved.

To address this limitation, we generated miR-27a KO, miR-27b KO, and miR-27a/b DKO mice, extending prior approaches that relied on antagomirs or deletion of entire miR-23/27/24 clusters^22, 24, 27, 33^. Our CRISPR-based strategy selectively excised the individual miR-27 hairpins while preserving transcription of adjacent miR-23 and miR-24 sequences, ensuring that observed phenotypes reflected specific loss of miR-27a or miR-27b rather than disruption of cluster architecture. This genetic framework allowed precise dissection of redundant and cooperative miRNA functions in adipose biology.

Using these models, we demonstrate that combined deletion of miR-27a and miR-27b produces the most robust enhancement of thermogenic function in subcutaneous and brown adipose tissues, with accompanying remodeling of gonadal fat toward a smaller, more multilocular adipocyte morphology. Cellular and tissue analyses support a cooperative repression model in which both miRNAs limit *Ucp1* induction and oxidative metabolism, thereby constraining thermogenic capacity. These findings align with established links between UCP1 activation, reduced lipid storage, and increased fatty acid oxidation^35, 36^, and indicate that miR-27a and miR-27b share overlapping regulatory targets that redundantly suppress thermogenic pathways.

Whole-body metabolic profiling further supports this model. Increased oxygen consumption and energy expenditure were observed in DKO mice during room temperature and cold exposure, whereas metabolic parameters at thermoneutrality remained similar to wild-type, indicating that miR-27 deficiency augments adaptive thermogenesis rather than basal metabolic rate. Improvements in glucose tolerance, fasting glycemia, adiposity, and adipose thermogenic gene expression under HFD conditions were most pronounced in the DKO, whereas single KOs produced only partial metabolic benefits. Together, these findings demonstrate that miR-27a and miR-27b cooperate to restrain thermogenic activation and promote energy storage, particularly under metabolic stress.

Comparison of our isoform-specific knockout models with prior cluster-deletion studies provides important biological insight. Global deletion of the miR-23b/27b/24-1 cluster impaired glucose tolerance and altered hepatic metabolic pathways, with both metabolomic and transcriptomic profiling indicating disrupted glycolysis and hepatic fuel utilization^26^. In contrast, skeletal muscle–specific deletion of the miR-23/27/24 clusters produced only subtle structural changes without affecting oxidative capacity, mitochondrial remodeling, or systemic metabolic performance^37^. These findings suggest that skeletal muscle is unlikely to mediate the systemic metabolic abnormalities observed in the global cluster knockout. Myeloid-specific deletion of the miR-23/27/24 clusters produced a paradoxical phenotype in which mice gained less weight on a high-fat diet yet exhibited worsened glucose and insulin tolerance^27^,. This outcome was attributed primarily to loss of miR-23 targeting of Eif4ebp2, a translational repressor that limits macrophage proliferation, resulting in a reduced pool of lipid-associated macrophages rather than to loss of miR-27 family members.

Taken together, these studies indicate that additional cluster-encoded miRNAs exert metabolic actions that differ from, and in some tissues oppose, those of the miR-27 family. As a result, whole-cluster deletion likely obscures the beneficial effects of miR-27 loss by simultaneously removing miRNAs that help maintain metabolic homeostasis. In contrast, our isoform-specific genetic strategy reveals a distinct gatekeeper role for miR-27a/b in restricting adipose thermogenesis and systemic metabolic flexibility, functions that cannot be resolved through cluster-level knockout approaches. Because nearly half of mammalian miRNAs are organized into polycistronic clusters, these findings demonstrate that miRNA clusters do not necessarily function as coherent regulatory modules and may contain antagonistic regulators whose individual contributions can be uncovered only through isoform-specific genetic dissection.

In a broader context, our work identifies miR-27a/b as regulatory nodes that influence the metabolic set point of adipose tissue. Their shared seed sequence and overlapping target networks suggest that they function as a coordinated post-transcriptional module integrating mitochondrial remodeling, nutrient status, and adipocyte fate decisions to determine whether adipose depots maintain a storage phenotype or transition toward a more oxidative, beige-like state. The depot-wide enhancement of thermogenic responsiveness observed following combined deletion further implies that miR-27 family members operate as gatekeepers of adipocyte recruitability rather than acting within a single fat subtype. These features highlight miR-27a/b as potential master regulators of adipose plasticity and systemic metabolic flexibility.

Beyond adipose biology, miR-27 family members have been implicated in diverse physiological and pathological processes, including skeletal remodeling^31^, angiogenesis and wound healing^38, 39^, fibrosis^40^, immune regulation^41, 42^ and cancer progression^43^. The genetically precise miR-27a, miR-27b, and double-knockout mouse strains generated here therefore represent valuable tools for interrogating miR-27 function across these additional biological systems under both physiological and disease conditions.

Future studies will focus on identifying additional miR-27 targets involved in regulating mitochondrial function, thermogenic signaling networks, and insulin resistance, and on defining how these regulatory pathways integrate across tissues to control whole-body energy balance, with the goal to advance miR-27-targeting strategies as potential therapeutic approaches for obesity and metabolic disease.

### Conclusions

This work illustrates how closely related miRNAs can act as an integrated regulatory module to shape adipose tissue plasticity and metabolic adaptability. Together, comparison of our isoform-specific knockout models with prior cluster-deletion studies indicates that miRNAs within the miR-23/27/24 loci exert opposing influences on metabolic regulation, such that the beneficial effects of miR-27 loss are masked when the entire cluster is deleted. The isoform-specific knockout models developed here provide a genetic framework to dissect how post-transcriptional networks coordinate mitochondrial remodeling, nutrient handling, and adipocyte fate decisions across tissues. Beyond establishing the role of the miR-27 family in thermogenic control, these models offer a powerful resource for defining miRNA-dependent mechanisms underlying metabolic disease and for testing targeted intervention strategies *in vivo*.

## Material and Methods

### Generation of miR-27a and miR-27b knockout mouse strains

To generate miR-27a and miR-27b knockout (KO) mice, CRISPR/Cas9 guide RNAs (gRNAs) were designed to target sequences flanking the hairpin regions of each microRNA within their native tri-miRNA clusters. For miR-27a, which resides within an intergenic tri-miRNA cluster (miR-23a/miR-27a/miR-24-2), gRNAs were positioned to excise the entire stem-loop structure of miR-27a while preserving the adjacent miR-23a and miR-24-2 sequences. For miR-27b, located within the intragenic cluster (miR-23b/miR-27b/miR-24-1) embedded in the *Aopep* and *Fancc* genes, gRNAs were designed within the intronic region to remove the miR-27b stem-loop without disrupting exon-intron boundaries or neighboring miRNAs.

CRISPR/Cas9 ribonucleoprotein complexes were delivered into C57BL/6J zygotes by pronuclear microinjection. Injected embryos were transferred into pseudo-pregnant surrogate females and carried to term. Founder mice were screened by PCR spanning the targeted deletion sites. Germline transmission was confirmed by outcrossing founders to wild-type (WT) C57BL/6J mice, followed by intercrossing of heterozygous offspring to generate homozygous KO animals. For genotyping, genomic DNA was isolated from toe biopsy samples and analyzed by PCR using primers flanking the deleted regions of miR-27a or miR-27b. PCR products of 313 bp for the miR-27a KO allele and 174 bp for the miR-27b KO allele were observed, whereas WT alleles yielded amplicons of 463 bp and 453 bp, respectively (Fig. 2C). PCR products were resolved by agarose gel electrophoresis and visualized using a ChemiDoc™ Imaging System (Bio-Rad Laboratories).

### Strain and Colony Maintenance

Homozygous C57BL/6 miR-27a^−^/^−^ (miR-27a KO) and miR-27b^−^/^−^ (miR-27b KO) mice were initially crossed with C57BL/6J wild-type (WT; +/+) mice to generate heterozygous offspring, which were subsequently intercrossed to produce experimental cohorts of male and female homozygous knockout and WT animals for each allele. To generate miR-27a/b double-knockout (DKO) mice, homozygous miR-27a KO mice were crossed with homozygous miR-27b KO mice to obtain double heterozygous progeny (miR-27a^+^/^−^; miR-27b^+^/^−^), which were intercrossed to generate male and female mice homozygous for both mutant alleles (miR-27a^−^/^−^; miR-27b^−^/^−^) along with WT controls. Routine colony maintenance was performed by intercrossing heterozygous miR-27a, miR-27b, and miR-27a/b mice of both sexes, with additional controlled breeding schemes including WT × WT, homozygous × homozygous, and heterozygous × heterozygous pairings to ensure continuous generation of all required genotypes and maintenance of stable experimental lines.

### Validation of miR-27a and miR-27b cluster expression

Mature miRNA levels were quantified by stem-loop reverse-transcription quantitative PCR (qPCR). For the miR-27a cluster, expression of miR-23a, miR-27a, and miR-24-2 was measured in WT and miR-27a KO mice. For the miR-27b cluster, expression of miR-23b, miR-27b, and miR-24-1 was analyzed in WT and miR-27b KO mice to assess the impact of miR-27 deletion on neighboring miRNAs within each native tri-miRNA cluster. To evaluate transcriptional integrity of the clusters following gene editing, PCR was performed using primers designed to amplify adjacent miRNA sequences flanking the deleted miR-27a or miR-27b loci. cDNA from WT, heterozygous, and homozygous KO samples was used as a template to verify stable expression of the remaining miRNAs within each cluster. For the intronic miR-27b cluster, expression and splicing of the host genes *Aopep* and *Fancc* were assessed by qPCR using primers spanning exon-exon junctions adjacent to the edited intron. These analyses confirmed that deletion of miR-27b did not alter host gene expression or transcript splicing.

### Assessment of breeding performance, fertility, and sex ratio

Independent breeding pairs of WT, miR-27a KO, miR-27b KO, and miR-27a/b DKO mice on a C57BL/6J background were established using age-matched males and females (8–16 weeks of age). For each breeding pair, the number of litters produced, total number of pups born, pup sex determined at birth, and the number of pups successfully weaned at postnatal day 21 were recorded. Breeding pairs that failed to produce offspring were also documented. All breeding performance data are summarized in Supplementary Table 1. Sex ratio was calculated as the proportion of male and female offspring relative to the total number of pups born. Fertility was defined as the percentage of breeding pairs that produced at least one litter out of the total number of pairs established. Mean litter size was determined as the average number of pups per successful litter. Comparisons of sex ratios and fertility rates among genotypes were performed using chi-square (χ^2^) tests to assess deviations from expected Mendelian ratios, while mean litter sizes were compared using one-way ANOVA with appropriate post hoc testing where applicable.

### Dietary intervention

For establishment of the high-fat diet (HFD) model, 3-week-old male miR-27a KO, miR-27b KO, and miR-27a/b DKO mice were placed on either a control diet (10% kcal from fat; Teklad TD.08806) or a high-fat diet (60% kcal from fat; Teklad TD.06414) for 15 weeks. Following the dietary intervention, glucose tolerance tests (GTT) and insulin tolerance tests (ITT) were performed. Mice were euthanized 48 hours after completion of ITT, and serum and tissue samples were collected for downstream analyses.

### Whole-body indirect calorimetry

Whole-body metabolic parameters were assessed using an automated indirect calorimetry system (Promethion Core, Sable Systems International, USA). Seven-week-old WT and DKO male mice were individually housed in rodent incubator under controlled environmental conditions with free access to standard chow and water. Prior to metabolic measurements, mice were acclimated for 6 days at thermoneutral temperature (28.5°C) in a rodent incubator. Following acclimation, mice were transferred to the Promethion Core Metabolic Cage System (Sable Systems International) for sequential temperature exposure. Mice were maintained at thermoneutral temperature (28.5°C) for 4 days, followed by room temperature (22°C) for 3 days, and then exposed to cold (6.5°C) for an additional 3 days. Throughout the experiment, metabolic parameters including whole-body energy expenditure, oxygen consumption (VO_2_), carbon dioxide production (VCO_2_), respiratory exchange ratio (RER), resting and active energy expenditure, physical activity, and food and water intake were continuously monitored and recorded using the Promethion Core system. The system was calibrated before each experimental run using standard reference gases (20.5% O_2_ and 0.5% CO_2_). Activity was quantified using infrared beam breaks for X, Y, and Z axes and expressed as total or ambulatory movements. EE and VO_2_ data were normalized to lean body mass (g) obtained by nuclear magnetic resonance (DXA, Faxitron Bioptics, LLC) prior to the experiment to account for body composition differences between genotypes.

### Glucose and insulin tolerance tests (GTT and ITT)

For GTT, mice maintained on either control or high-fat diets were fasted for 12 h. Baseline fasting blood glucose levels were obtained prior to intraperitoneal injection of glucose (2 g/kg body weight). Blood glucose was then measured at 15, 30, 60, 90, and 120 min post-injection. For ITT, mice were fasted for 6 h before baseline blood glucose measurement and intraperitoneal administration of insulin (0.5 IU/kg body weight). Blood glucose levels were subsequently recorded at 15, 30, 60, 90, and 120 min following insulin injection. All blood glucose measurements were performed using an Accu-Chek Guide Me glucometer (Roche Diagnostics).

### *In vitro* adipocyte culture and differentiation

Preadipocytes were isolated from interscapular BAT and inguinal WAT of 8-week-old male WT, miR-27a KO, miR-27b KO, and miR-27a/b DKO mice. Dissected tissues were digested in tissue lysis buffer containing 0.123 M NaCl, 1.3 mM CaCl_2_, 5 mM glucose, 100 mM HEPES, 4% (v/v) bovine serum albumin (BSA; fraction V), and 0.1% (w/v) Collagenase P at 37 °C for 30-40 min with constant rocking. Following complete digestion, the lysate was filtered through a 70-µm cell strainer with an equal volume of autoMACS® Running Buffer to reduce cell clumping. The filtrate was centrifuged at 1200 × g for 5 min, and the resulting pellet was washed with 1× PBS and centrifuged again at 500 × g for 5 min. The final cell pellet was resuspended in growth medium consisting of DMEM supplemented with 10% fetal bovine serum (FBS), 1× GlutaMAX, and 1× antibiotic–antimycotic solution, and plated onto 10-cm or 15-cm culture dishes according to cell yield. For differentiation, preadipocytes were seeded into 12-well or 24-well plates in growth medium. Upon reaching full confluence, cells were incubated for 3 days in differentiation medium composed of DMEM supplemented with 1.7 µM insulin, 1 µM triiodothyronine (T3), 5 µM rosiglitazone, 10 µM SB431542, 500 µM IBMX, 10 µM dexamethasone, 125 µM indomethacin, and 50 µg/mL L-ascorbic acid 2-phosphate (AA2P). After 3 days, the medium was replaced with maintenance medium consisting of DMEM containing 1.7 µM insulin, 1 µM T3, 5 µM rosiglitazone, 10 µM SB431542, and 50 µg/mL AA2P, and cells were incubated for an additional 3 days. Thereafter, cultures were maintained in growth medium until day 10 of differentiation.

### Oil Red O Staining

Preadipocytes were isolated from inguinal WAT of WT and miR-27a/b DKO mice and cultured under standard adipogenic differentiation conditions as described above. On day 10 of differentiation, cells were washed once with phosphate-buffered saline (PBS) and fixed with 10% NBF for 15 min at room temperature. Fixed cells were incubated with 0.3% Oil Red O working solution for 15 min, followed by five washes with distilled water to remove excess dye. Stained cultures were imaged using an EVOS M5000 Imaging System (Thermo Fisher Scientific). For quantitative analysis, Oil Red O dye was eluted by incubating stained cells with 100% isopropanol for 10 min on an orbital shaker at room temperature. Absorbance of the extracted dye was measured in duplicate at 500 nm using a GloMax luminometer (Promega).

### Mitochondrial DNA copy number analysis

Mitochondrial DNA (mtDNA) content was quantified by qPCR using primers specific for the mitochondrial gene mt-Nd1 and a single-copy nuclear reference gene (Rplp0 or B2m). Absolute mtDNA copy number was determined using standard curves generated from serial dilutions of plasmids containing the corresponding amplicons of known sequence length. The theoretical copy number of each dilution was calculated from plasmid size and DNA concentration using Avogadro’s number. Copy numbers for both mitochondrial and nuclear targets were interpolated from their respective standard curves, and mtDNA content was normalized to nuclear DNA levels to calculate relative mtDNA copy number per cell. All assays were performed according to established protocols^44-46^.

### RNA isolation and quantitative PCR

Total RNA from cells and tissues was extracted using the miRNeasy Micro Kit (Qiagen) according to the manufacturer’s instructions. RNA concentration and purity were assessed by NanoDrop™ 2000 spectrophotometry (Thermo Fisher Scientific), with acceptable 260/280 absorbance ratios ranging from 1.9 to 2.0. For mRNA analysis, 300 ng of total RNA was reverse transcribed using the qScript cDNA Synthesis Kit (Quanta Biosciences). Quantitative PCR (qPCR) was performed using AzuraView™ GreenFast qPCR Blue Mix LR (Azura Genomics) on a CFX Opus 384 Real-Time PCR System (Bio-Rad) with gene-specific primers (Supplementary Table 2). For microRNA analysis, 20 ng of RNA was reverse transcribed using the TaqMan™ MicroRNA Reverse Transcription Kit (Thermo Fisher Scientific) with miRNA-specific TaqMan™ primers for miR-27a and miR-27b. miRNA qPCR was performed using TaqMan™ Fast Universal PCR Master Mix on the same Bio-Rad platform with miRNA-specific TaqMan™ probes. β-Actin and 18S rRNA were used as endogenous controls for normalization of mRNA and miRNA expression, respectively. Relative expression levels were calculated using the ΔΔCt method.

### Protein isolation and Western blot analysis

Protein lysates were prepared from cells and tissues using RIPA lysis buffer (Sigma-Aldrich) supplemented with protease and phosphatase inhibitor cocktails (Sigma-Aldrich). Equal amounts of total protein (20 µg per sample) were separated by SDS-PAGE and transferred to PVDF membranes for immunoblotting. Membranes were blocked and incubated with a primary antibody against UCP1 (Cell Signaling Technology; 1:1,000 dilution), followed by incubation with an HRP-conjugated secondary antibody (Cell Signaling Technology; 1:5,000 dilution). Immunoreactive bands were detected using Clarity™ and Clarity Max™ ECL Western blotting substrates (Bio-Rad). β-Tubulin or β-Actin served as loading controls for normalization. Densitometric analysis of band intensities was performed using ImageJ software, with UCP1 signals normalized to the corresponding loading controls after background subtraction.

### Tissue Collection, Processing, and Histological Analysis

At the conclusion of the high-fat diet (HFD) study, mice were euthanized and blood was collected by cardiac puncture. Serum was isolated by centrifugation at 1,300 × g for 10 min at 4 °C, aliquoted, and stored at −80 °C until subsequent analysis. Adipose tissue samples designated for RNA extraction were homogenized in QIAzol using a bead-beating homogenizer and stored at −80 °C until further processing. Protein samples were snap-frozen in liquid nitrogen and stored at −80 °C for downstream analyses. For histological evaluation, brown adipose tissue (BAT), inguinal white adipose tissue (iWAT), and epididymal white adipose tissue (eWAT) were harvested from wild-type (WT) and miR-27a/b double-knockout (DKO) mice maintained on control or high-fat diets at the completion of dietary intervention. Tissues were fixed in 10% neutral-buffered formalin (NBF) for 48 h at 4 °C, dehydrated through graded ethanol beginning at 70%, and embedded in paraffin. Paraffin blocks were sectioned at 5 µm thickness using a rotary microtome and mounted onto positively charged glass slides. Routine hematoxylin and eosin (H&E) staining was performed, and stained sections were imaged by bright-field microscopy using an EVOS M5000 Imaging System (Thermo Fisher Scientific).

### Heat Production Assay via Microcalorimetry

Heat production from differentiated beige adipocytes or freshly isolated adipose tissue was measured using high-resolution isothermal microcalorimetry on the calScreener™ system (Symcel, Sweden). For in vitro measurements, beige preadipocytes were expanded and differentiated in DMEM medium (as described above) on Geltrex-coated calWell™ insert plates (Symcel), then transferred into screw-capped titanium vials for analysis. The calPlate™ (48-well format) was pre-equilibrated between two thermal zones for 30 min, followed by a stabilization period of 15-30 min after final positioning. Continuous heat output was recorded using calView™ software. For ex vivo analysis, interscapular brown adipose tissue and inguinal white adipose tissue were collected from 8-week-old male wild-type (WT), miR-27a knockout (KO), miR-27b KO, and miR-27a/b double-knockout (DKO) mice. Tissue biopsies were obtained from standardized anatomical locations within each fat pad, placed into calWells containing 200 μL DMEM supplemented with 1% bovine serum albumin (BSA), 1× GlutaMAX (Gibco), and 1× antibiotic–antimycotic solution (Gibco), and normalized to tissue weight prior to analysis. Samples were incubated in the calScreener system for 16 h. All measurements were normalized to internal reference wells containing medium alone.

### Statistical Analysis

All experiments were performed with biological triplicates or greater, and data are presented as mean ± standard error of the mean (SEM). Statistical comparisons between two groups were performed using unpaired Student’s t-tests, while comparisons among multiple groups were analyzed by one-way or two-way analysis of variance (ANOVA) followed by appropriate post hoc testing as applicable. Statistical significance was defined as p < 0.05.

### Animal Ethics

All animal studies were conducted in compliance with the National Institutes of Health Guide for the Care and Use of Laboratory Animals and were approved by the Institutional Animal Care and Use Committee (IACUC; protocol #2203) at the MaineHealth Institute for Research. Mice were housed in a pathogen-free barrier facility with ad libitum access to food and water and maintained on a 12-hour light/dark cycle with food and water provided ad libitum. All efforts were made to minimize animal suffering and reduce animal numbers whenever possible.

## Data Availability

All data supporting the findings of this study are provided within the manuscript and accompanying Supplementary Information. Raw and processed datasets from RNA and microRNA qPCR analyses, microcalorimetry experiments, and metabolic cage studies are available from the corresponding author upon reasonable request.

## Acknowledgements

This work was supported by National Institute of Diabetes and Digestive and Kidney Diseases (NIDDK) award R01DK124261 (A. Brown) and National Institutes of Health COBRE award P20GM121301 (A. Brown, L. Liaw, and C. J. Rosen). This research utilized core facilities supported by NIH award U54GM115516 (C. J. Rosen, Principal Investigator). The authors gratefully acknowledge the substantial experimental contributions provided by the Genome Modification, Physiology, Flow Cytometry, and Histology Cores at the MaineHealth Institute for Research.

---

**Supplementary Table 1.**
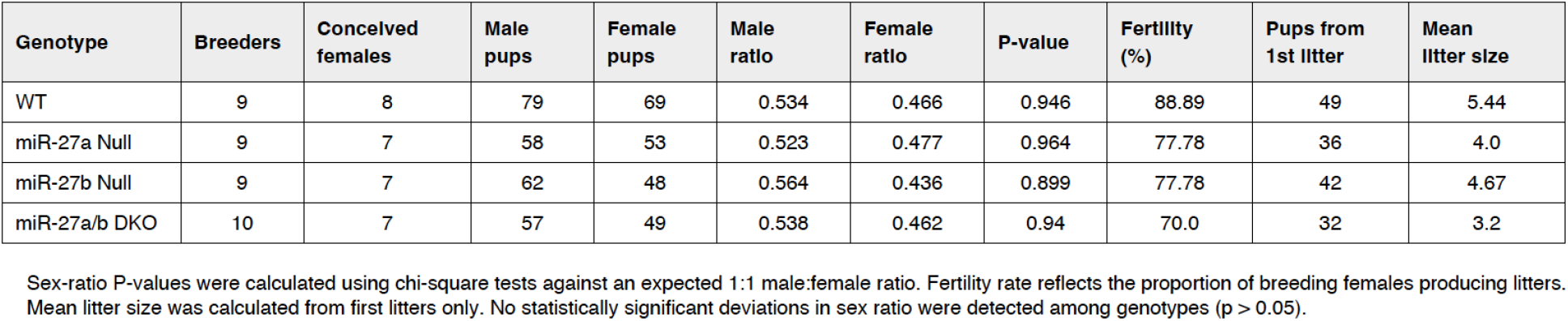
Breeding performance and sex distribution of miR-27 knockout lines.

| Genotype | Breeders | Conceived females | Male pups | Female pups | Male ratio | Female ratio | P-value | Fertility (%) | Pups from 1st litter | Mean litter size |
| --- | --- | --- | --- | --- | --- | --- | --- | --- | --- | --- |
| WT | 9 | 8 | 79 | 69 | 0.534 | 0.466 | 0.946 | 88.89 | 49 | 5.44 |
| miR-27a Null | 9 | 7 | 58 | 53 | 0.523 | 0.477 | 0.964 | 77.78 | 36 | 4.0 |
| miR-27b Null | 9 | 7 | 62 | 48 | 0.564 | 0.436 | 0.899 | 77.78 | 42 | 4.67 |
| miR-27a/b DKO | 10 | 7 | 57 | 49 | 0.538 | 0.462 | 0.94 | 70.0 | 32 | 3.2 |

Sex-ratio P-values were calculated using chi-square tests against an expected 1:1 male:female ratio. Fertility rate reflects the proportion of breeding females producing litters. Mean litter size was calculated from first litters only. No statistically significant deviations in sex ratio were detected among genotypes ( $p > 0.05$ ).

**Supplementary Table 2.**
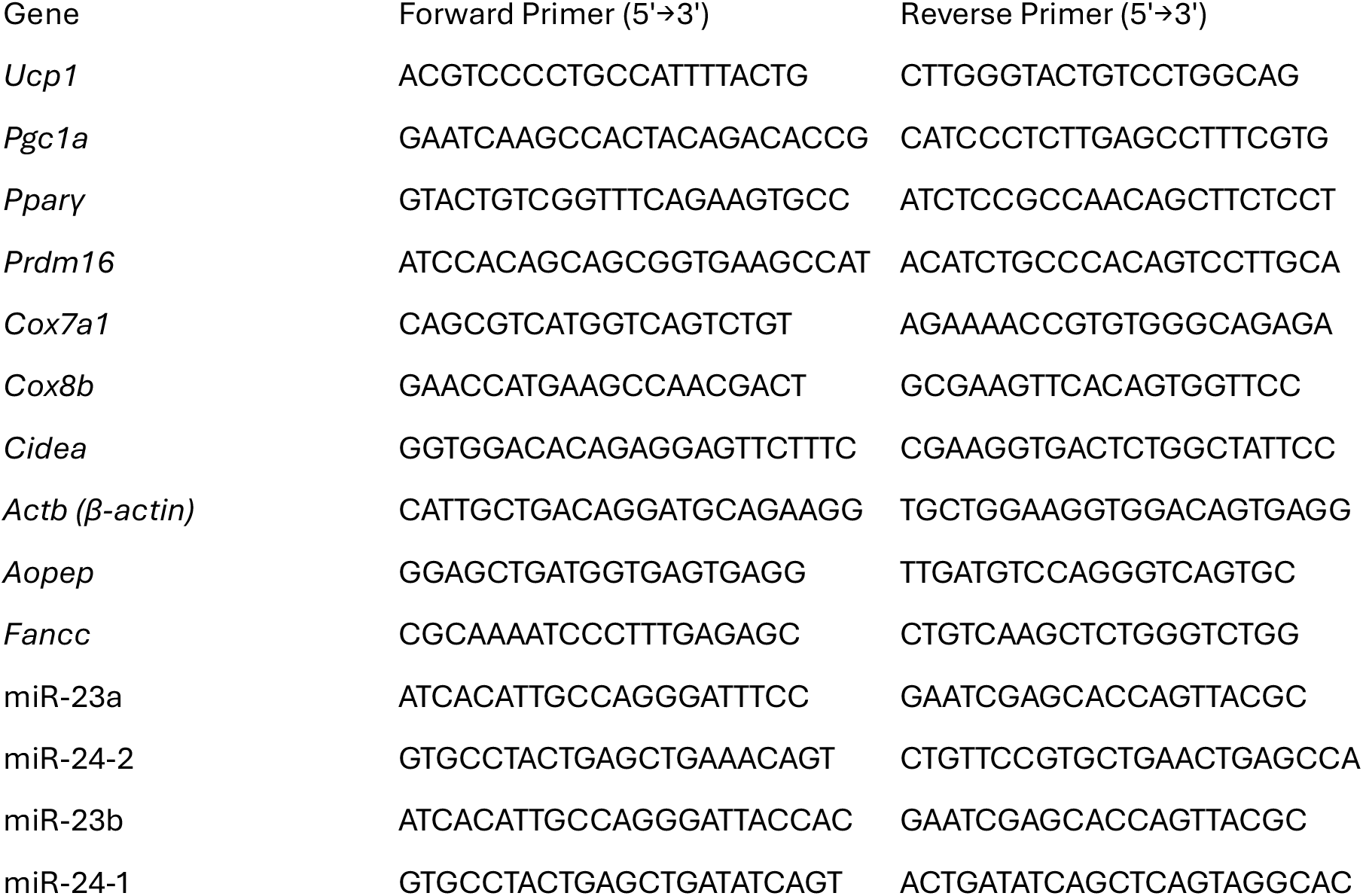
Primer sequences used for quantitative PCR.

## Notes

### Competing Interest Statement

The authors have declared no competing interest.

### Summary of Updates

Additional data added, additional authors added, complete rewrite of the manuscript including title change.

