## Supplemental figures for "Isoform-Specific Control of Adipose Thermogenesis by the miR-27 Family Reveals Antagonism within a Polycistronic miRNA Cluster"

### Supplementary Figure 1

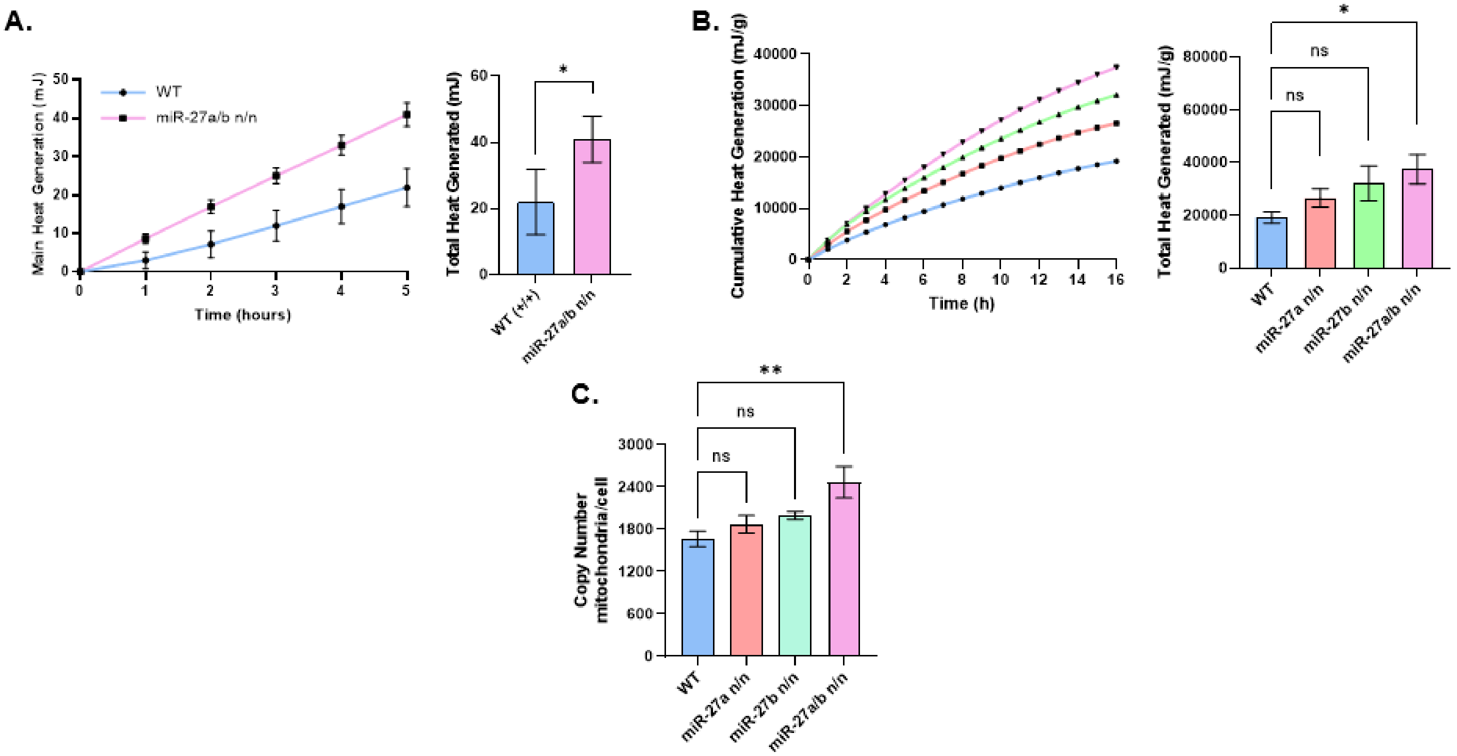


**Supplementary Figure 1. Thermogenic function and mitochondrial abundance in brown adipocytes and brown adipose tissue. (A)** Isothermal microcalorimetric analysis of heat production from cultured brown adipocytes differentiated from stromal vascular fraction isolated from brown adipose tissue (BAT) of WT and miR-27a/b DKO mice (n = 5). Left panel shows cumulative heat production over time; right panel displays quantification of total heat output derived from time-course measurements. **(B)** Isothermal microcalorimetric analysis of heat generation from BAT explants isolated from 8-week-old WT, miR-27a knockout (KO), miR-27b KO, and miR-27a/b DKO mice (n = 7). Left panel shows cumulative heat production over time; right panel shows quantification of total heat output. **(C)** Mitochondrial DNA copy number analysis in BAT isolated from 8-week-old WT, miR-27a KO, miR-27b KO, and miR-27a/b DKO mice (n = 4). Data are presented as mean ± SEM. Statistical significance is indicated as *p < 0.05, **p < 0.01, and ****p < 0.0001.

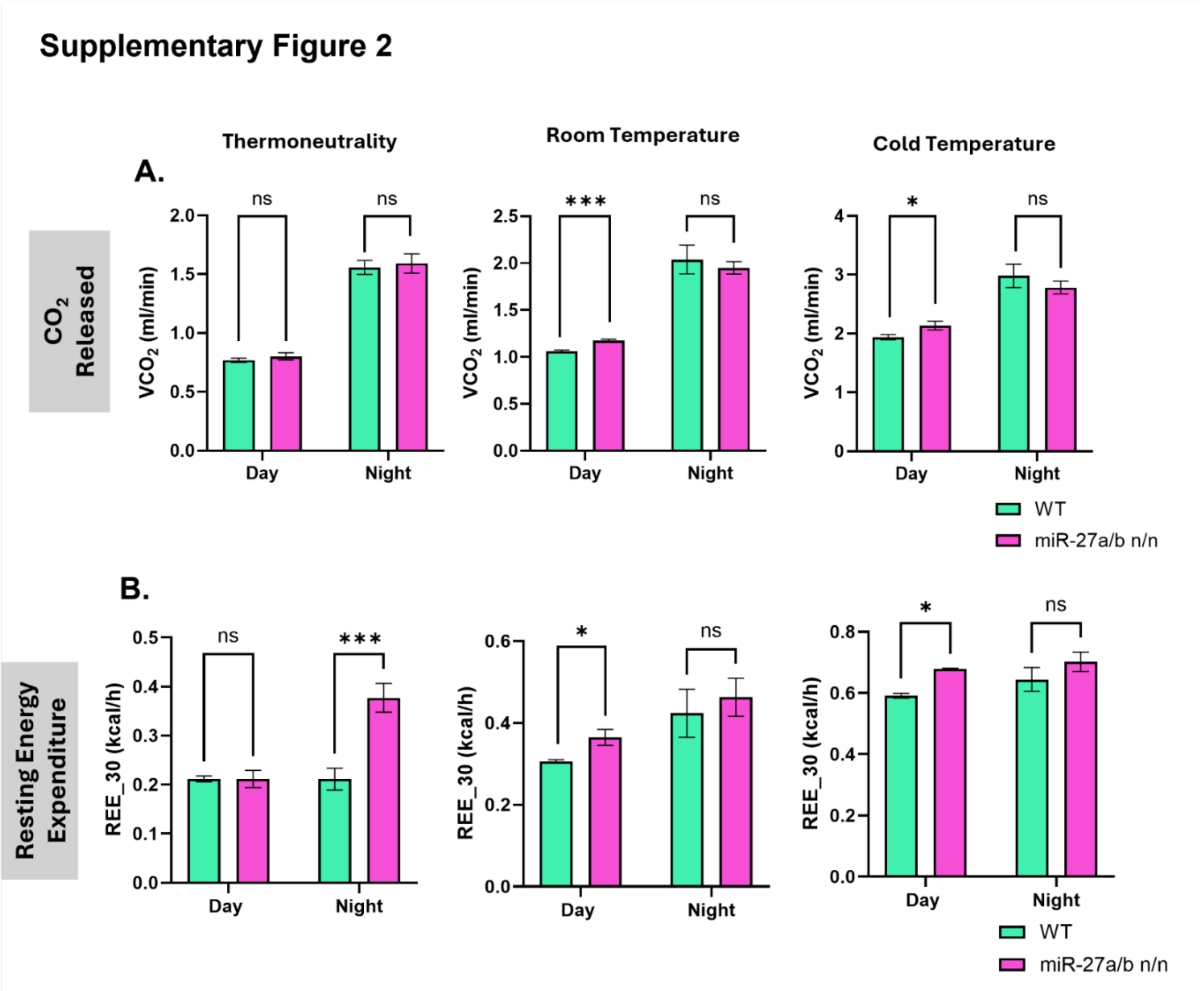


**Supplementary Figure 2. Metabolic cage assessment of respiratory output and resting energy expenditure in WT and miR-27a/b DKO mice. (A)** Carbon dioxide production ($VCO_2$) measured in wild-type WT and miR-27a/b DKO mice under thermoneutral (28 °C), room temperature (22 °C), and cold (6.5 °C) conditions. **(B)** Mean resting energy expenditure calculated during the lowest 30-minute activity period ($REE_{30}$) in WT and DKO mice across the same temperature conditions. Data are presented as mean ± SEM. Statistical significance is indicated as *$p < 0.05$ and ***$p < 0.001$.

### Supplementary Figure 3

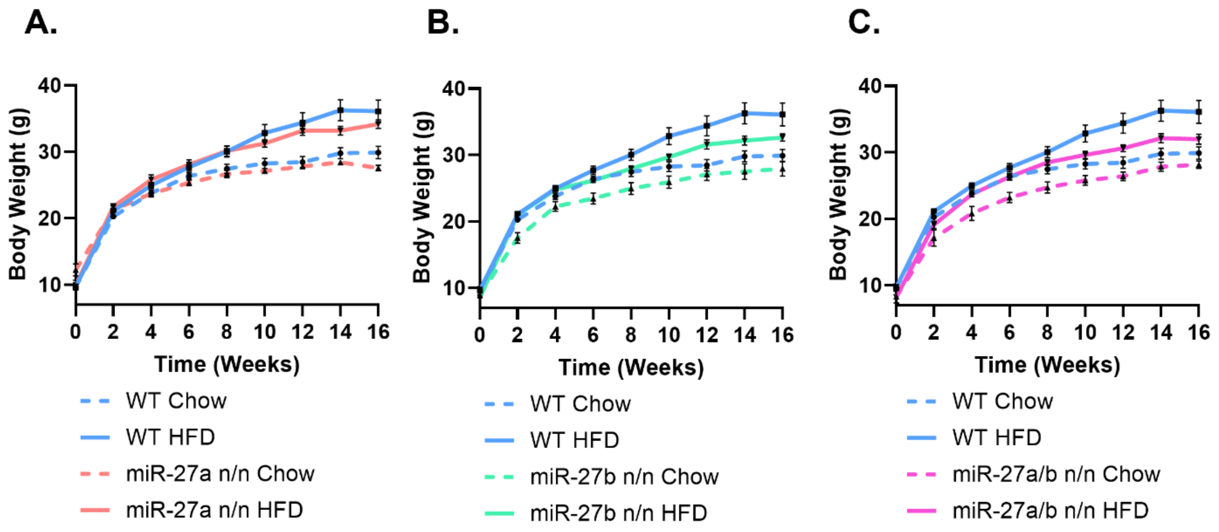


**Supplementary Figure 3. Body weight trajectories during chow and high-fat diet feeding in miR-27 knockout mice. (A-C)** Longitudinal body weight measurements of male miR-27a KO **(A)**, miR-27b KO **(B)**, and miR-27a/b DKO **(C)** mice maintained on standard chow or high-fat diet (HFD), shown in comparison with WT controls (n = 9 per group). Data are presented as mean ± SEM.
